# Single-type neuron transcriptomics uncovers cell-type-specific regulators of host defense in *Caenorhabditis elegans*

**DOI:** 10.64898/2026.05.08.723815

**Authors:** Phillip Wibisono, Yiyong Liu, Jingru Sun

## Abstract

Understanding how the nervous system regulates immune responses requires insight into how individual neurons respond to infection. In *Caenorhabditis elegans*, sensory neurons such as ASH play important roles in modulating innate immunity; however, the molecular mechanisms operating within these neurons remain poorly defined. Previous transcriptomic studies have relied on whole-animal RNA sequencing, which lacks the cellular resolution needed to detect neuron-specific signaling programs. Here, we performed single-type neuron transcriptomic profiling to characterize gene expression in ASH neurons from infected and uninfected animals. We found that ASH neurons undergo extensive transcriptional remodeling in response to pathogen exposure, with enrichment of genes associated with G protein-coupled receptor signaling, neuropeptide activity, and sensory transduction. Notably, we identified genes, including *zig-3* and *F36F2*.*8*, that are strongly induced in ASH neurons but are not detected in whole-animal transcriptomic datasets. Functional analysis using ASH-specific RNA interference demonstrated that knockdown of either gene significantly reduces survival during *Pseudomonas aeruginosa* infection, whereas whole-animal RNAi produces no detectable phenotype. Together, these findings reveal that cell-type-resolved transcriptomics can uncover functionally important regulators of host defense that are masked in bulk analyses and provide new insight into neuron-intrinsic mechanisms of neuroimmune regulation.

## INTRODUCTION

The ability to sense and respond to environmental challenges is essential for organismal survival, and the nervous system plays a central role in coordinating these responses. In addition to regulating behavior, increasing evidence indicates that neural circuits actively modulate physiological processes, including innate immunity (1-7). Across species, the nervous system integrates external sensory cues with internal signaling pathways to fine-tune immune responses, thereby maintaining homeostasis during infection (1, 3, 4, 8-13). Despite these advances, a fundamental question remains unresolved: how do individual neurons detect pathogenic stimuli and translate these signals into molecular outputs that regulate host defense?

The nematode *Caenorhabditis elegans* provides a powerful model to address this question due to its well-defined nervous system and conserved immune pathways (10). Previous studies have demonstrated that specific sensory neurons regulate immune responses through neuroendocrine signaling mechanisms (4, 5, 8, 14-20). In particular, the amphid sensory neuron ASH has emerged as a key regulator of innate immunity. Prior work from our group and others showed that ASH neurons regulate immune activation through signaling pathways involving the G protein-coupled receptor OCTR-1 and downstream modulation of conserved immune pathways, including the p38 MAPK and unfolded protein response (UPR) pathways (4, 5, 21). These findings established a functional link between sensory perception and immune regulation. However, the molecular mechanisms operating within ASH neurons, specifically how these neurons respond transcriptionally to pathogen exposure, remain unknown.

A major obstacle to resolving neuron-intrinsic mechanisms of immune regulation has been the reliance on whole-animal transcriptomic approaches. Bulk RNA sequencing provides a global view of gene expression changes during infection but lacks the cellular resolution necessary to distinguish neuron-specific responses. In *C. elegans*, ASH neurons consist of a single bilateral pair (22), representing an extremely small fraction of the total somatic cell population (two ASH neurons out of approximately 959 cells in the hermaphrodite). Consequently, transcriptional changes occurring within these neurons are highly diluted in whole-worm lysates and are therefore difficult to detect using conventional approaches. As a result, neuron-specific regulators of immune signaling may remain obscured in bulk transcriptomic analyses.

To overcome these limitations, we employed a single-type neuron RNA sequencing approach described by Spencer *et al*. (23) to selectively profile gene expression in ASH neurons. By isolating ASH neurons from infected and uninfected animals and performing low-input transcriptomic analysis, we directly captured neuron-intrinsic transcriptional responses to pathogen exposure. This strategy enables the identification of signaling molecules and pathways that are specifically activated within ASH neurons and that may mediate neural control of innate immunity.

In this study, we show that ASH neurons undergo extensive transcriptional remodeling in response to infection and identify candidate genes that are selectively induced in this neuron type but not detected in whole-animal transcriptomic datasets. Functional analyses further demonstrate that these genes contribute to host survival in a neuron-specific manner. Together, our findings reveal previously unrecognized components of neuroimmune regulation and highlight the importance of cell-type-resolved approaches for uncovering molecular mechanisms that are obscured in bulk analyses.

## RESULTS

### Overview of the single-type neuron RNA-seq workflow

Previously, we have demonstrated that in *C. elegans*, the amphid ASH sensory neurons suppress the innate immune response to pathogen infection (4). However, the molecular details of signal transduction in these neurons remain unknown. To gain insights into such neural regulations, we employed single-type neuron RNA-seq to profile gene expression in these cells with or without pathogen infection. Figure 1 depicts the workflow of our method. Briefly, we generated a transgenic *C. elegans* strain (JRS111) expressing GFP specifically in ASH neurons using the Q repressible binary expression system (24) (Figure 2A). The synchronized JRS111 animals were exposed to the pathogenic bacterium *Pseudomonas aeruginosa* strain PA14 or the non-pathogenic control *Escherichia coli* strain OP50 for 4 hours as previously described (4). Following exposure, infected and uninfected animals were collected and subjected to transient treatment with SDS-DTT to pre-sensitize worm cuticle, and then digested with pronase E to remove the cuticle, followed by pipetting to release individual cells (25, 26). After that, we obtained cell suspension that contained dissociated single cells. The cells were then sorted through our Beckman-Coulter CytoFlex SRT cell sorter with a 100-micron diameter nozzle to enrich GFP-labeled ASH neurons (Figure 2B). Non-fluorescent cells from wild-type N2 animals were used as a reference to exclude autofluorescent cells. cDNA was synthesized directly from neuronal cells by using SMART-Seq Ultra Low Input RNA Kit v4 (Takara Bio). cDNA library was constructed using Nextera XT DNA Library Preparation Kit (Illumina), followed by sequencing on an Illumina NovaSeq 6000 sequencer. The resulting sequence data were subjected to alignment and differential gene expression analysis to identify regulated genes in the infected worms relative to uninfected controls, following our established protocols (8). We also performed gene ontology (GO) analysis and function assays to examine gene functions in worm defense against infection. Below we describe our method and results in detail.

**Figure 1.**
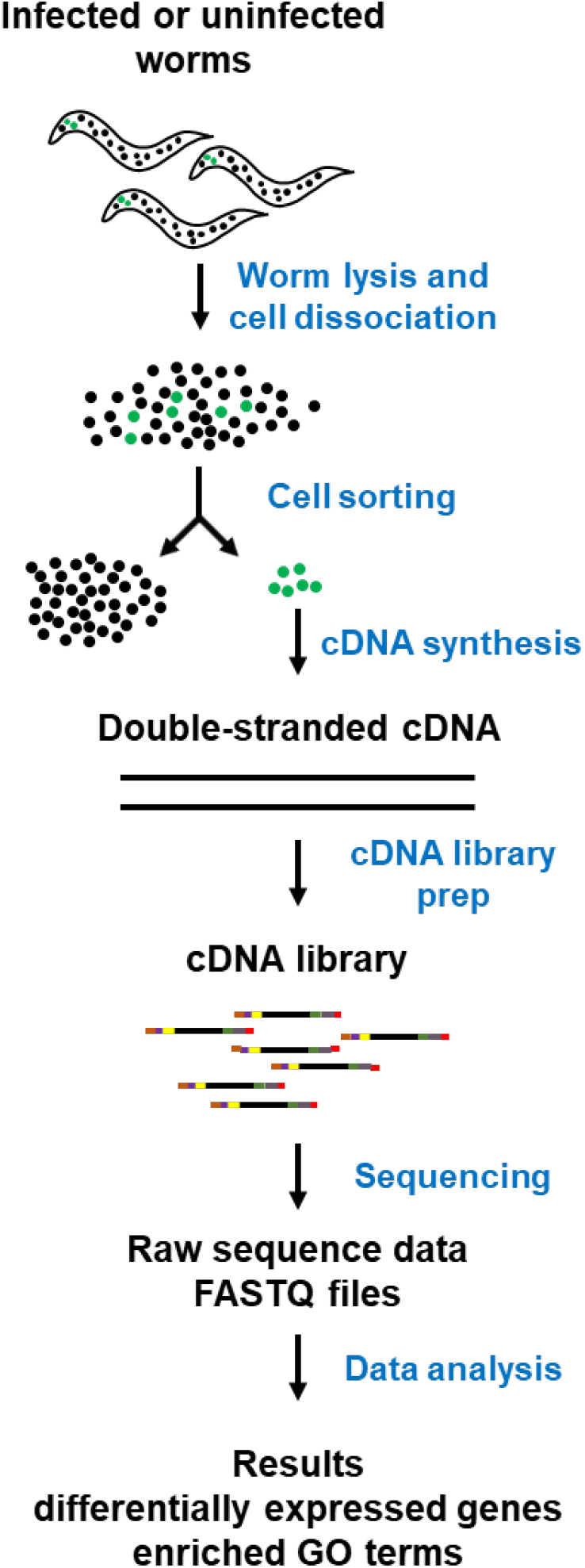
Overview of single-type neuron RNA sequencing in ASH neurons. Schematic of the experimental pipeline. Transgenic *C. elegans* expressing GFP in ASH neurons were exposed to *Pseudomonas aeruginosa* PA14 or *Escherichia coli* OP50, followed by dissociation into single cells. GFP-labeled ASH neurons were isolated by fluorescence-activated cell sorting (FACS), and RNA was processed for low-input sequencing. Differential gene expression, gene ontology analysis, and functional assays were performed to identify neuron-specific regulators of host defense.

**Figure 2.**
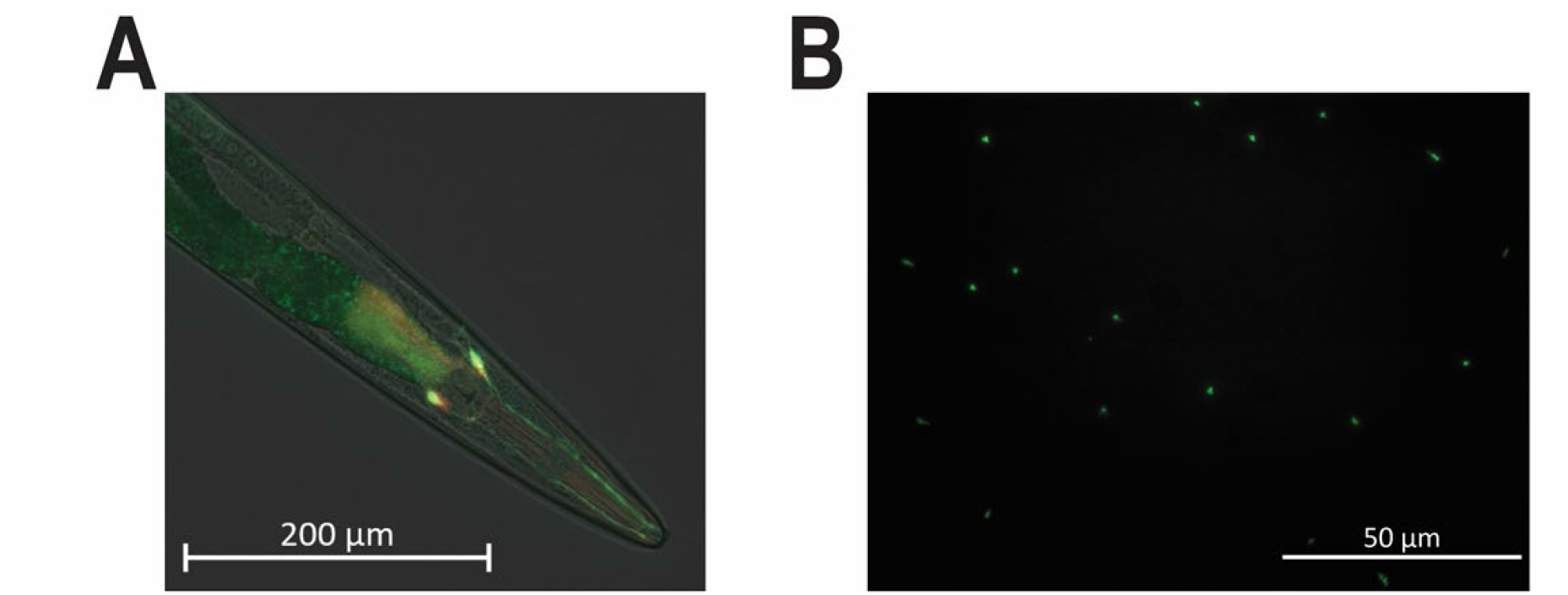
Isolation and enrichment of ASH neurons for single-type neuron RNA sequencing. (A) Representative fluorescence image of the transgenic *C. elegans* strain JRS111 expressing GFP specifically in ASH neurons. (B) Isolation and enrichment of ASH neurons by FACS.

### Evaluation of sequence data quality from single-type neuron RNA-seq

In this study, we collected ASH neurons from both uninfected animals (four biological replicates) and *P. aeruginosa*-infected animals (five biological replicates) for library preparation and sequencing. Approximately 20 million paired-end reads with a length of 100 bp were generated for each sample, which are sufficient for downstream gene expression quantification and differential expression analysis (27).

We first evaluated the quality of sequence data by examining several key parameters, including quality scores of base sequences, alignment rate, alignment distribution, read coverage over the position of transcripts, and duplicates. To this end, we compared these parameters between our single-type neuron RNA-seq and standard bulky RNA-seq. In our previous study, we performed standard bulky RNA-seq on wild-type *N2* animals with or without *P. aeruginosa* infection (8). Here, we compared the quality of sequence data between the uninfected samples from the two types of RNA-seq. Both RNA-seq generated high-quality base sequences with almost all bases sequenced above Phred score 30 (Q30). For alignment rate, 84.6% of the reads from the single-type neuron RNA-seq were aligned to the reference genome (Fig. 3A). While this rate is acceptable (28), it is significantly lower than the alignment rate (99.5%) of the standard RNA-seq (Fig. 3A). Among the aligned reads in the single-type neuron RNA-seq, 86.31% corresponded to coding sequences, 10.72% to untranslated regions (UTRs), 0.4% to intronic sequences, and 2.58% to intergenic sequences, all of which were comparable to the percentages observed in the standard RNA-seq (Fig. 3B). While the coverage across transcripts from 5’ to 3’ was balanced in the standard samples, such coverage in the single-type neuron RNA-seq was slightly biased towards the 3’ end (Fig. 3C), indicating that the 5’ ends of the RNA molecules in the latter were either partially degraded or were not sufficiently transcribed during cDNA synthesis or both. The single-type neuron samples also had more duplicates than the standard RNA-seq (57.88% vs. 27.08%) (Fig. 3D). Overall, the sequence data quality of the single-type neuron RNA-seq is slightly worse than that of our standard bulky RNA-seq but is still better than the data quality of many low-input and bulky RNA-seq studies (29, 30).

**Figure 3.**
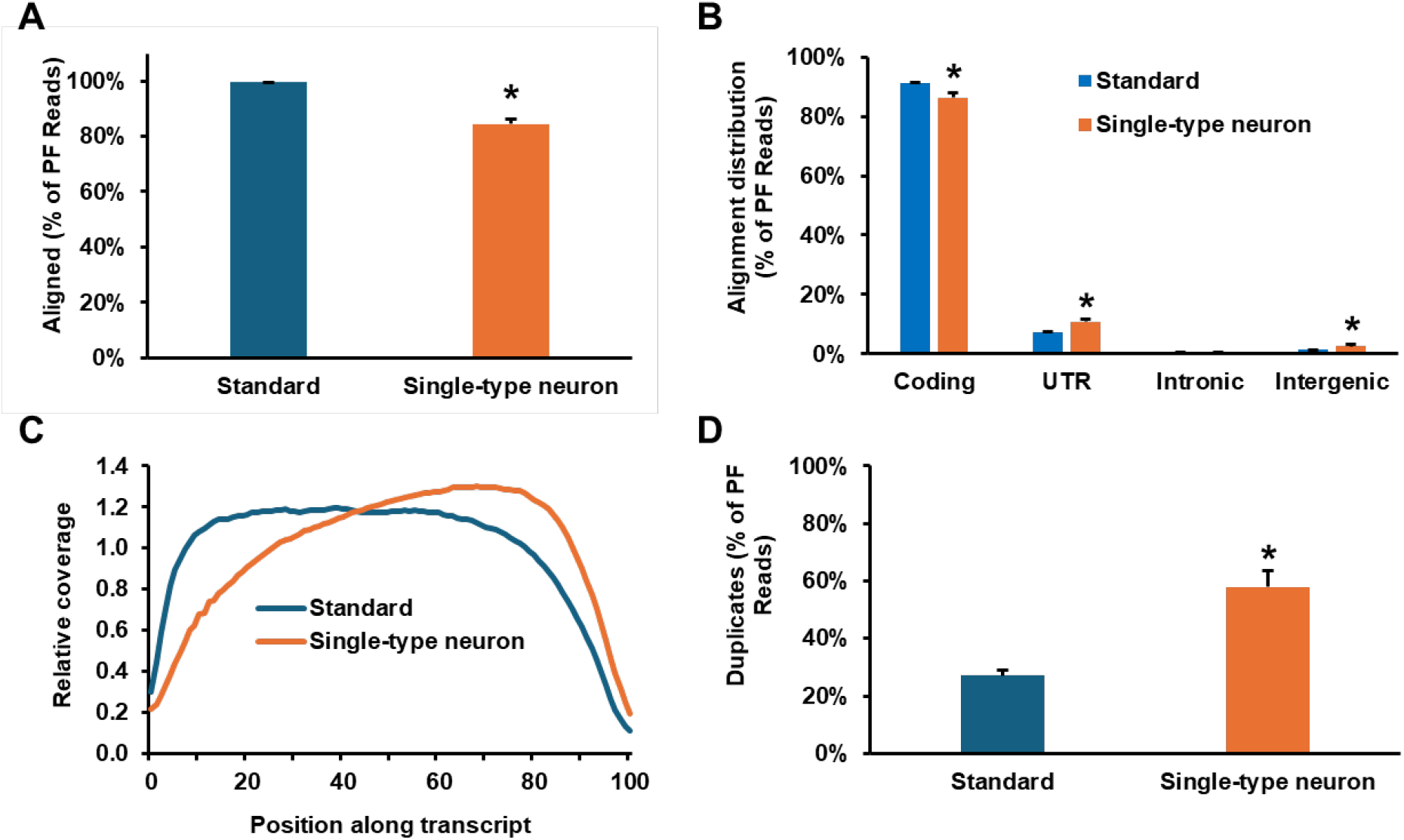
Quality metrics of single-type neuron and standard RNA-seq libraries. (A) Alignment rates, expressed as the percentage of reads mapped to the C. elegans reference genome relative to total passing-filter (PF) reads. Each bar in the histogram represents the mean value of four libraries. Error bars indicate standard deviation. P < 0.05 (asterisk) denotes a statistically significant difference between library types. (B) Genomic distribution of aligned reads, shown as the percentage of PF reads mapping to coding regions, untranslated regions (UTRs), intronic regions, and intergenic regions. Each bar in the histogram represents the mean value of four libraries, with standard deviation indicated by error bars. P < 0.05 (asterisk) denotes a statistically significant difference between library types. (C) Transcript coverage profiles across gene bodies. Transcripts were normalized and divided into 1,000 equal bins from 5′ to 3′ ends. Each curve represents the average of four libraries. (D) Duplicate read rates, calculated as the percentage of duplicate reads among total PF reads. Each bar in the histogram represents the mean value of four libraries, with standard deviation indicated by error bars. P < 0.05 (asterisk) denotes a statistically significant difference between library types.

### ASH neurons undergo extensive transcriptional remodeling upon infection

In our ASH single-type neuron RNA-seq from uninfected animals, a total of 13,656 genes/transcripts were identified and quantified with a false discovery rate (FDR) of 5%. Comparing these genes to those detected by the standard bulk RNA-seq identified 2,967 genes enriched in ASH neurons. Of these 2,967 genes, 112 overlaps with 147 previously reported ASH-enriched genes from a single-cell RNA-seq study (Figure 4A) (31). This overlap is highly significant (p < 6.63 × 10^−57^), providing strong validation of the specificity of our ASH single-type neuron RNA-seq and also substantially expanding the number of ASH-enriched genes.

**Figure 4.**
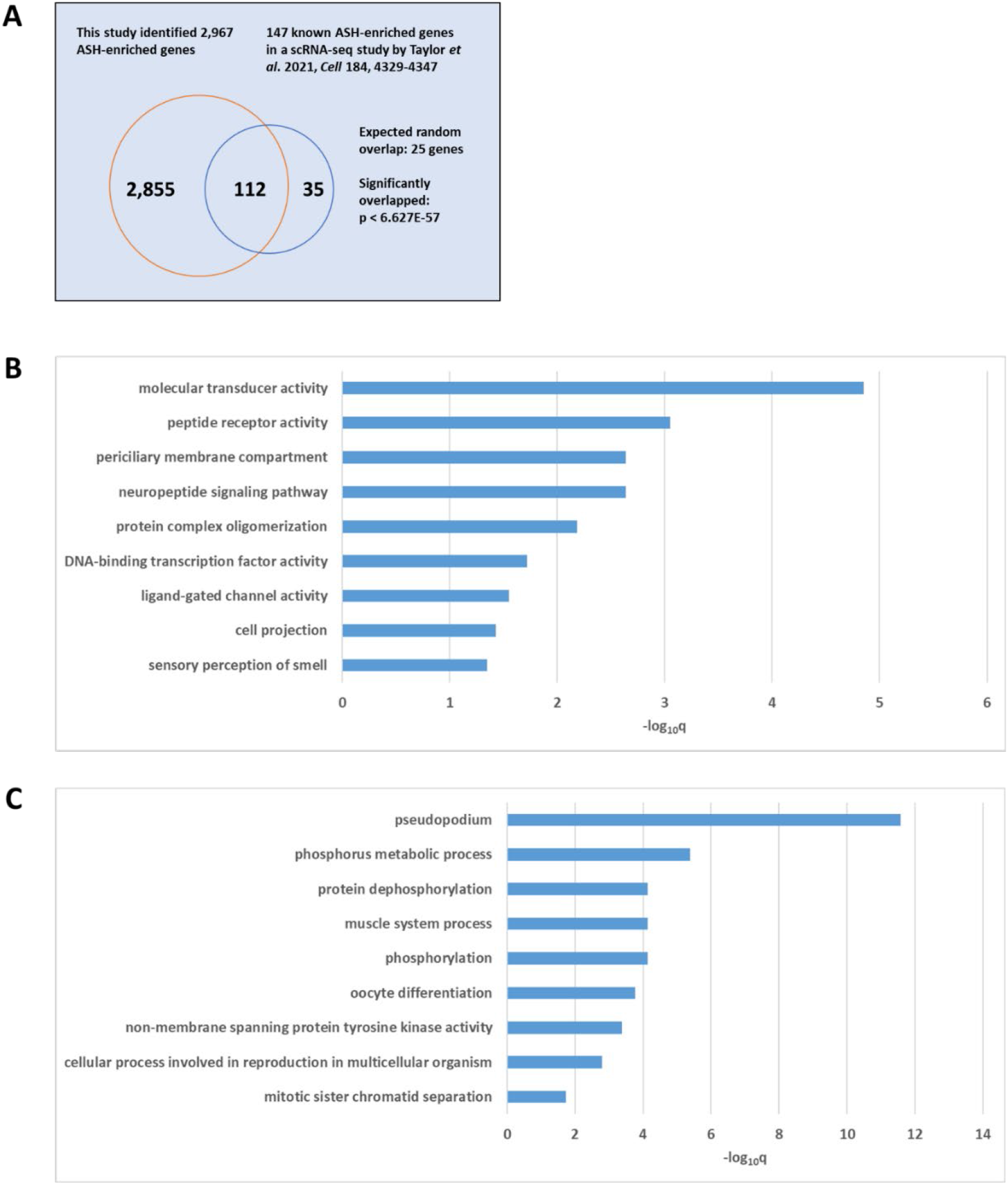
Transcriptional profiling and functional enrichment of ASH neurons. (A) Overlap between ASH-enriched genes identified in this study and previously reported ASH-specific genes from a single-cell RNA-seq dataset (Taylor et al., 2021). A total of 2,967 ASH-enriched genes were identified in this study, with 112 genes overlapping with 147 previously reported ASH-enriched genes. The overlap is highly significant (p < 6.63 × 10^−57^), indicating strong validation of dataset specificity. (B) Gene ontology (GO) enrichment analysis of genes upregulated in ASH neurons following Pseudomonas aeruginosa infection, highlighting enrichment of pathways related to molecular transducer activity, peptide receptor activity, and neuropeptide signaling. (C) GO enrichment analysis of genes downregulated in ASH neurons upon infection, showing enrichment of pathways associated with phosphorus metabolic processes and protein dephosphorylation.

To examine how pathogen infection affects gene expression in ASH neurons, we compared the expression profiles of ASH neurons isolated from *P. aeruginosa*-infected animals with those isolated from uninfected control animals. This comparison identified 189 genes upregulated and 853 genes downregulated by at least twofold. A gene ontology (GO) analysis of the 189 upregulated genes revealed significant enrichment of pathways related to molecular transducer activity, peptide receptor activity, and neuropeptide signaling (Figure 4B). A GO analysis of the 853 downregulated genes revealed the downregulation of phosphorus metabolic process and protein dephosphorylation (Figure 4C). These results indicate that ASH neurons actively respond to infection by modulating signaling pathways associated with environmental sensing and intercellular communication.

### Single-type neuron transcriptomics reveals functionally important neuroimmune regulators masked in whole-animal analyses

To determine whether ASH-specific transcriptomics can uncover genes that are not detected in conventional bulk analyses, we compared our single-type neuron RNA-seq dataset with previously generated whole-worm RNA-seq data from infected wild-type animals. Although bulk RNA-seq identified numerous infection-responsive genes, some transcripts that were strongly induced in ASH neurons were not detected as differentially expressed at the whole-animal level. This observation suggests that infection-induced transcriptional responses occurring within a small subset of neurons can be masked in whole-animal analyses, highlighting the importance of cell-type–resolved approaches for identifying neuron-intrinsic signaling molecules.

To determine whether these ASH-enriched genes contribute to host defense, we performed neuron-specific RNA interference (RNAi) targeting *zig-3* and *F36F2*.*8*, two genes that were robustly upregulation in ASH neurons following pathogen exposure but were absent from prior bulk RNA-seq datasets. For ASH-specific RNAi, we utilized the transgenic strain JRS112, which was generated by MosSCI-mediated single-copy insertion of a cassette expressing *sid-1* under the control of the ASH-specific *sra-6* promoter (*sra-6p::GFP::SL2::3xsynSID-1*) at the ttTI10882 genomic locus. This design enables selective RNAi sensitivity in ASH neurons, which are otherwise RNAi-resistant, while leaving other tissues unaffected. The construct also includes a hygromycin resistance marker for selection of transgenic animals.

To restrict RNAi activity to ASH neurons, JRS112 was crossed into the *sid-1(qt9)* mutant background to generate the strain JRS113, thereby eliminating systemic RNAi and confining gene knockdown to ASH neurons expressing *sid-1*. JRS113 animals subjected to ASH-specific knockdown of either *zig-3, F36F2*.*8*, or empty vector (RNAi control) were exposed to *Pseudomonas aeruginosa* PA14, and animal survival was monitored over time as previously described (4, 8, 16, 17, 21, 32-35). Knockdown of either *zig-3* or *F36F2*.*8* resulted in significantly increased susceptibility to infection compared to control animals treated with empty vector RNAi, indicating that both genes are required for normal host survival during pathogen infection (Figure 5A).

**Figure 5.**
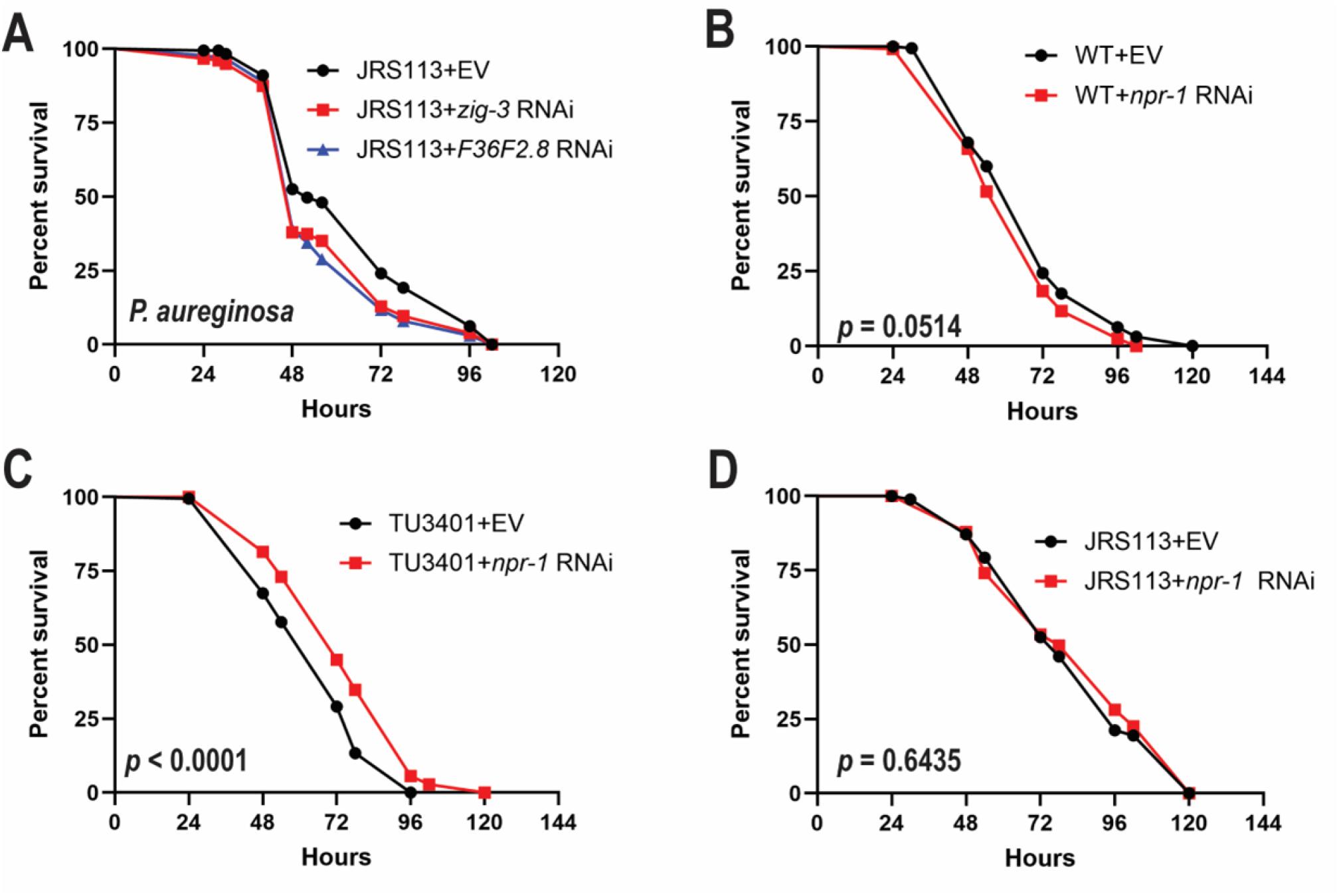
Functional validation of ASH-enriched genes identified by single-type neuron RNA-seq. (A) Survival of JRS113 animals with ASH-specific RNAi knockdown of *zig-3* or *F36F2*.*8* following *Pseudomonas aeruginosa* PA14 infection. Knockdown of either gene significantly reduced survival compared to animals treated with empty vector (EV) RNAi controls (*zig-3* RNAi, p = 0.0048; *F36F2*.*8* RNAi, p = 0.0004). (B) Survival of wild-type animals subjected to *npr-1* RNAi following PA14 infection (p = 0.0514). (C) Survival of pan-neuronal RNAi-sensitive TU3401 animals subjected to *npr-1* RNAi following PA14 infection (p < 0.0001). (D) Survival of JRS113 animals subjected to *npr-1* RNAi following PA14 infection (p = 0.6435).

To validate the specificity of our RNAi system, we performed control experiments using wild-type (WT), pan-neuronal RNAi-sensitive strain TU3401, and the ASH-specific RNAi strain JRS113. RNAi clones targeting *npr-1* gene were used as controls. *npr-1* gene is expressed in multiple sensory neurons, including ASH (14, 36, 37). As expected, *npr-1* RNAi led to reduced survival in the pan-neuronal RNAi-sensitive TU3401 strain but did not produce a significant decreased survival phenotype in WT or ASH-specific JRS113 animals (Figure 5B-D).

Together, these results demonstrate that RNAi-mediated gene knockdown occurs in a cell-type-specific manner in these genetic backgrounds and validate the use of JRS113 for selective manipulation of gene expression in ASH neurons. More broadly, our findings establish that single-type neuron transcriptomics can uncover biologically important regulators of host defense that are obscured in whole-animal analyses, revealing a previously hidden layer of neuroimmune regulation.

## DISCUSSION

In this study, we demonstrate that single-type neuron transcriptomics can reveal functionally important regulators of host defense that are not detectable using conventional whole-animal approaches. By focusing on a defined neuronal subtype, we provide direct evidence that ASH sensory neurons undergo extensive transcriptional reprogramming in response to pathogen infection and actively contribute to host immune regulation.

A key finding of this work is the identification of *zig-3* and *F36F2*.*8* as infection-induced genes in ASH neurons that are required for host survival during *Pseudomonas aeruginosa* infection. Notably, these genes were not detected in previous whole-animal RNA-seq datasets, highlighting a major limitation of bulk transcriptomic approaches. The observation that ASH-specific RNAi knockdown results in increased susceptibility to infection, whereas whole-animal RNAi does not produce a phenotype, indicates that the function of these genes is spatially restricted and dependent on cell-type-specific context. These findings provide direct functional evidence that neuron-intrinsic transcriptional responses can have organism-level consequences for host defense.

Our results support a model in which ASH neurons are not merely passive regulators but actively respond to pathogen exposure through dynamic transcriptional remodeling. The enrichment of genes associated with peptide receptor activity, neuropeptide signaling pathway, and molecular transducer activity suggests that ASH neurons may integrate environmental signals and modulate immune responses through intercellular communication pathways. These findings are consistent with previous studies demonstrating that neuronal signaling pathways, including OCTR-1-mediated regulation of the unfolded protein response and MAPK pathways, play central roles in coordinating immunity in *C. elegans* (4, 5, 21).

More broadly, this study highlights the importance of cell-type-resolved transcriptomic approaches for uncovering hidden layers of biological regulation. As demonstrated here, genes with critical functional roles can remain undetected in whole-organism analyses due to signal dilution but become apparent when examined at the level of individual cell types. This principle is likely applicable beyond *C. elegans*, particularly in complex systems where rare or specialized cells play disproportionate roles in physiological regulation.

Despite these advances, several questions remain. The molecular mechanisms by which *zig-3* and *F36F2*.*8* mediate neuroimmune signaling are not yet defined, and it will be important in future studies to determine how these genes interact with known immune pathways such as PMK-1 and the unfolded protein response. In addition, extending this approach to other neuronal subtypes may reveal additional layers of neural regulation of immunity.

In conclusion, our study establishes single-type neuron transcriptomics as a powerful strategy for dissecting neuron-intrinsic responses to infection and provides new insight into the molecular basis of neuroimmune regulation. By revealing regulators that are masked in bulk analyses, this work uncovers a previously hidden dimension of host defense and lays the foundation for future mechanistic studies of neural control of immunity.

## Resource availability

### Lead contact

Information and request regarding resources and reagents should be directed to and will be fulfilled by the lead contact, Jingru Sun

### Materials availability

The *C. elegans* strains and recombinant DNA generated in this study will be shared upon request, but we may require payment to cover shipment and completion of a Material Transfer Agreement for possible commercial applications.

## Acknowledgments

We would like to thank the *Caenorhabditis* Genetics Center (CGC) for providing worm strains used in this study. The CGC is funded by the NIH Office of Research Infrastructure Programs (P40 OD010440). This work was financially supported by the NIH, United States (R35GM158259 to Y.L. and R35GM124678 to J.S.). The funder had no role in the study design, data collection and interpretation, or the decision to submit the work for publication.

## Author contributions

P. W. and Y. L. designed and performed experiments and analyzed data. J. S. designed experiments and analyzed data. J. S., Y. L., and P. W. wrote the manuscript.

## Declaration of interests

The authors declare no competing interests.

## STAR Methods

- Key Resource Table
- Resource Availability
  - Lead contact
  - Materials availability
  - Data and code availability
- Experimental Model and Subject Details
  - Nematode Strains
  - Bacteria Strains
- Method Details
  - Plasmid construction and transgenic animal generation
  - Neuron isolation and flow cytometry
  - RNA sequencing
  - RNAi interference
  - Survival Assay
  - RNA isolation
  - Quantitative real-time PCR (qRT-PCR)
  - Quantification and statistical analysis

## KEY RESOURCES TABLE

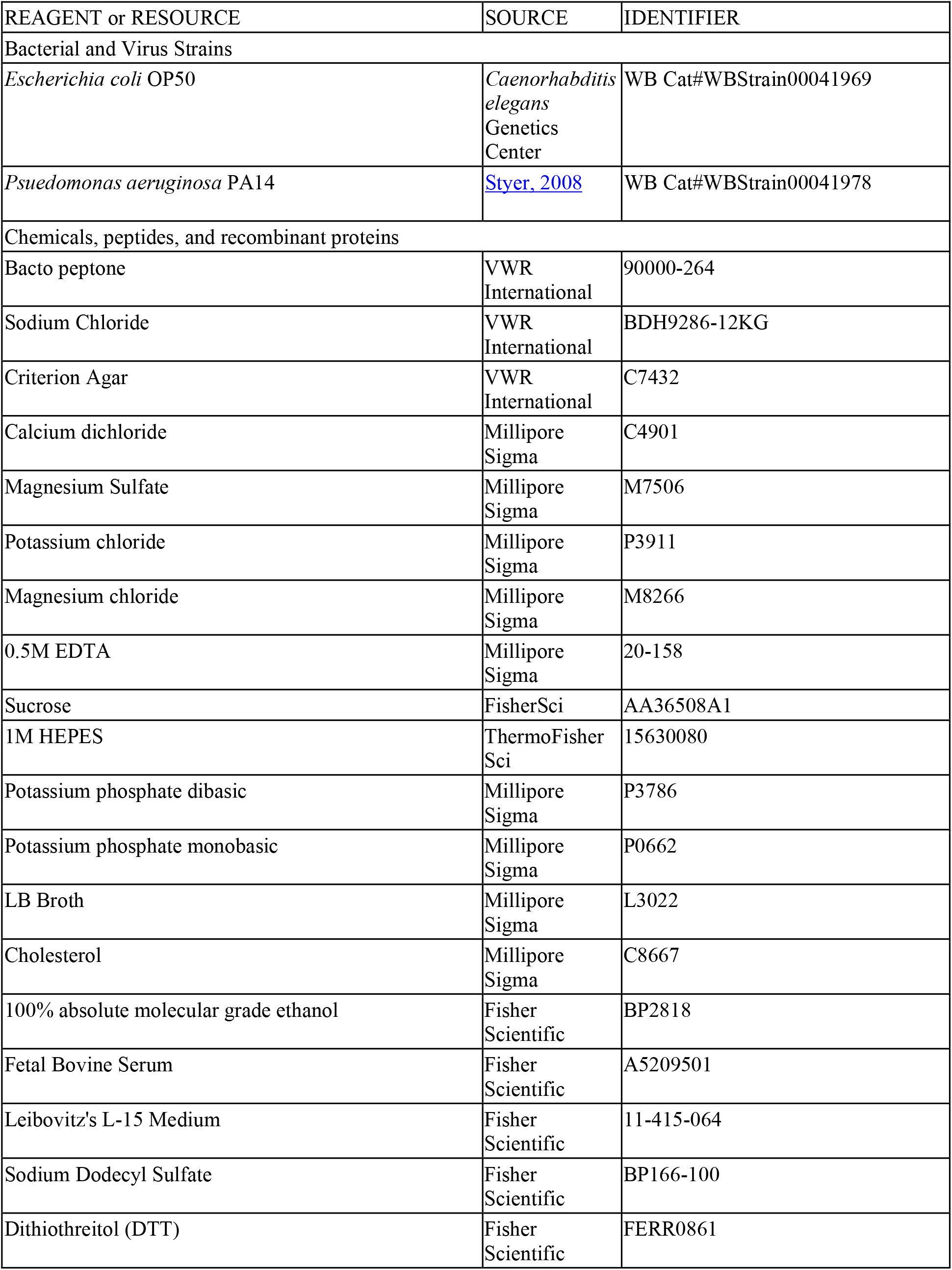
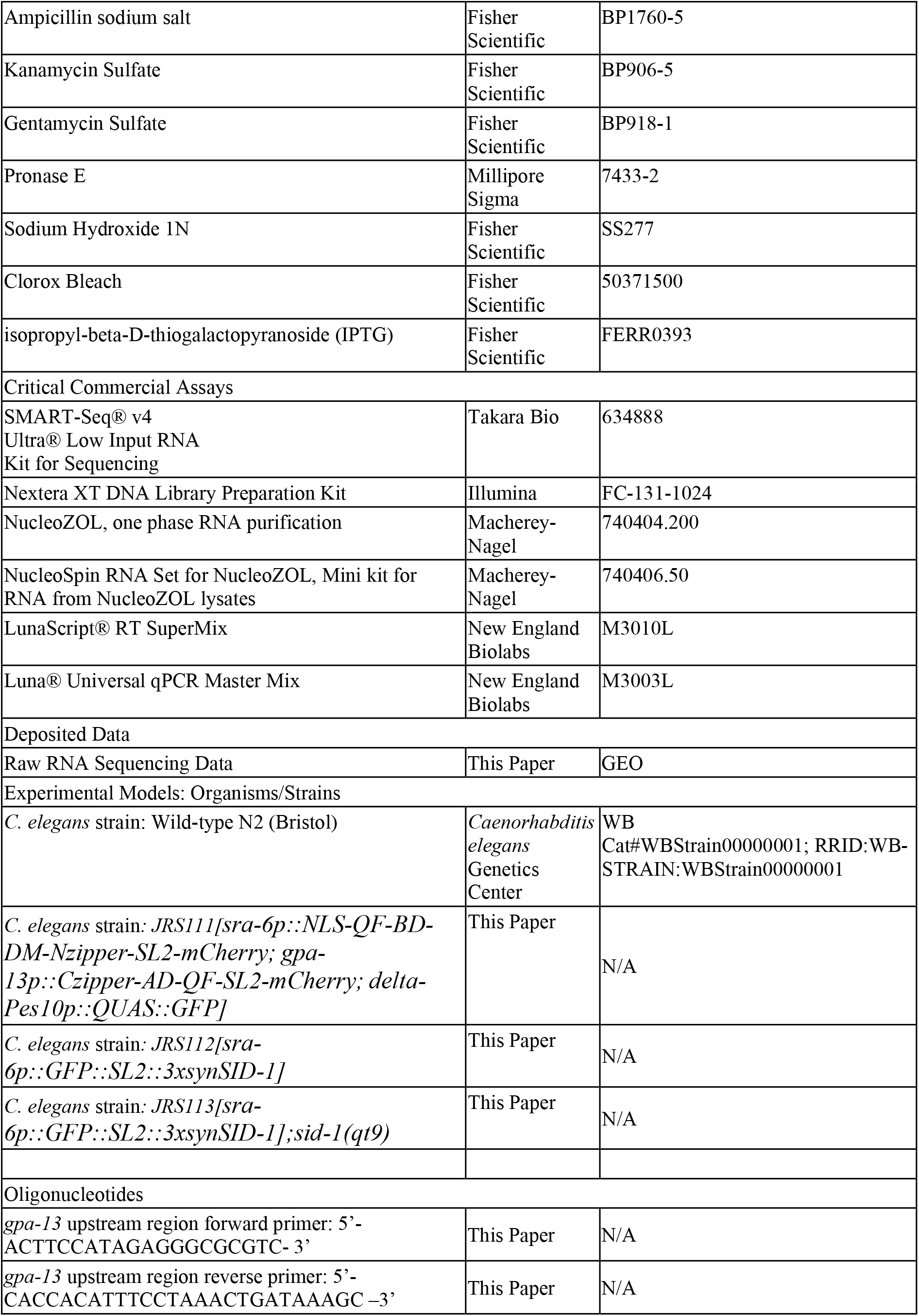
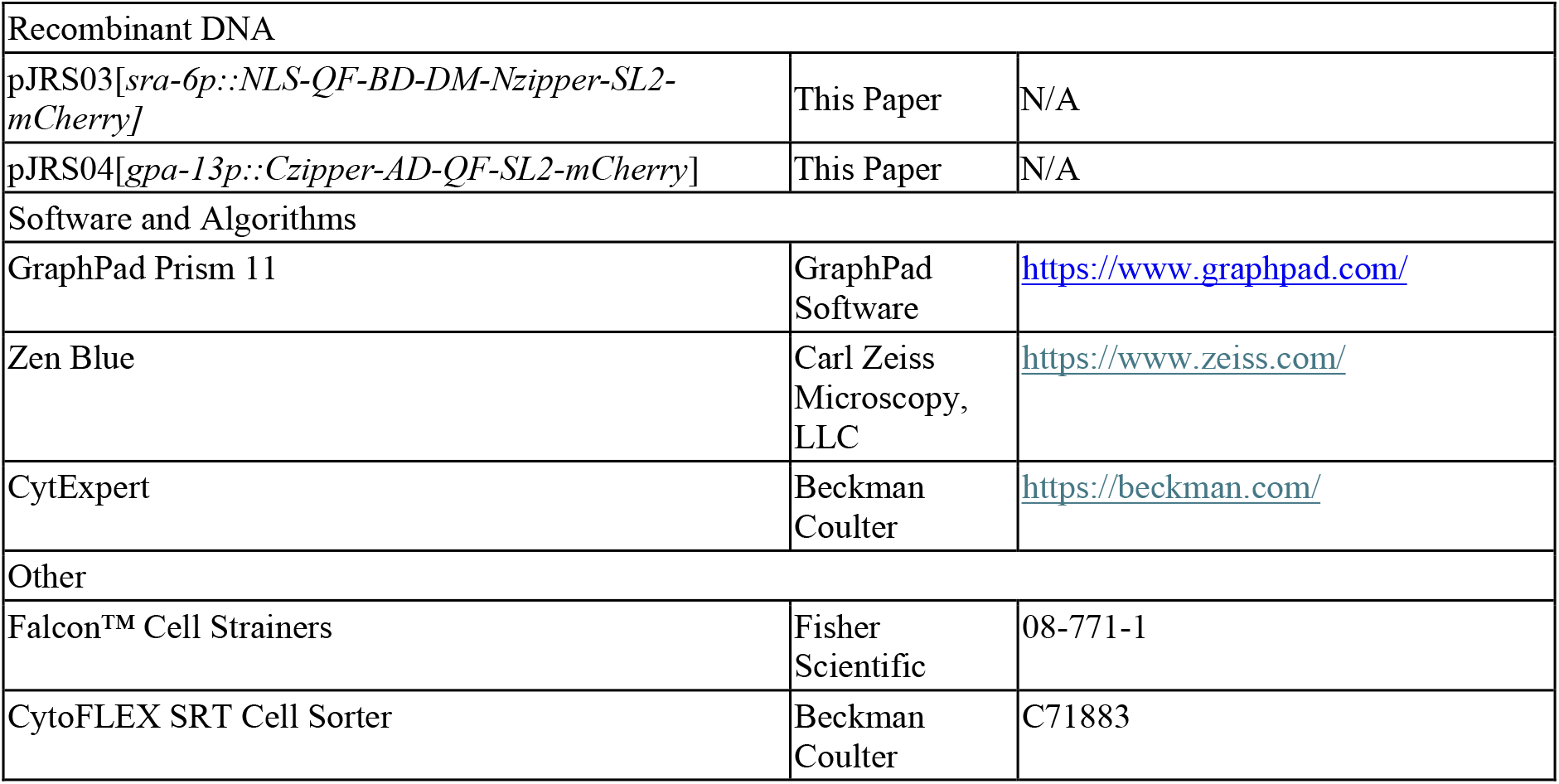

### Experimental model and subject details

#### Nematode strains

The following *C. elegans* strains were maintained as hermaphrodites at 20°C, grown on Nematode Growth Media (NGM) with 30 units/mL nystatin, and fed *E. coli* OP50. The wild-type animal strain was *C. elegans* Bristol *N2*. All animals used for this study were synchronized 65-hour old young adults unless otherwise noted in the specific methodology.

#### Bacterial strains

The following bacterial strains were grown using standard conditions: *Escherichia coli* OP50, *Pseudomonas aeruginosa* strain PA14.

## METHOD DETAILS

### Plasmid construction and transgenic animal generation

The upstream DNA fragment (3,793bp) upstream of *sra-6* was digested using the restriction enzymes NotI and AscI from the plasmid pTM139 and inserted into XW54 (Kang Shen, Addgene #65839) to generate pJRS03.

A genomic DNA fragment (2,241bp) upstream of *gpa-13* was amplified by PCR from a mixture of various stages of wild-type *C. elegans* using the following primers: 5’ - ACTTCCATAGAGGGCGCGTC-3’ and 5’ – CACCACATTTCCTAAACTGATAAAGC-3’. The PCR products were first cloned into pCR-TOPO 2.1(Invitrogen, Cat#K450002) before digestion with restriction enzymes *SphI* and *BamHI*. The resulting DNA fragment was ligated into XW52 vector (Kang Shen, Addgene #65838) digested with the same restriction enzymes to make pJRS04.

pJRS03, pJRS04, and XW12 (Kang Shen, Addgene#65384) were injected into wild-type animals at the following concentrations to generate JRS111: pJRS03 at 50ng/µL, pJRS04 at 50ng/µL, and XW12 at 10ng/µL.

JRS112 strain which expressed the gene *sid-1* in ASH neurons was generated by InVivo Biosystems by inserting the following cassette; [*sra-6p::GFP::SL2::3xsynSID-1::sra-6u; rps-0p::HygR::unc-54u*] using MosSCI at the ttTI10882 insertion site. A 2000bp upstream region of *sra-6* is placed ahead of *sid-1* cDNA with synthetic introns and the *sra-6* 3’UTR sequence. A hygromycin resistance gene was also inserted as a selective marker for successful integration.

JRS113 was generated by crossing JRS112 with *sid-1(qt9)* using standard genetic techniques and is resistant to RNA interference with exception of ASH neurons.

### Neuron isolation and flow cytometry

JRS113 animals expressing GFP solely in ASH neurons grew at 20°C on NGM seeded with *E. coli* OP50. Animals were synchronized by lysing adult hermaphrodite with a solution of sodium hydroxide and blech (ration 5:2). The freed embryos were washed and were allowed to develop for 22hrs in S-basal liquid media at 20°C. Synchronized L1 larval animals were transferred to modified NGM plates seeded with *E. coli* OP50 and allowed to grow for 48 hours at 20°C until the L4 larval stage. After 48 hours, approximately 100 thousand synchronized L4 larval animals were transferred onto either modified NGM plates seeded with *E. coli* OP50 or *P. aeruginosa* PA14 and incubated at 25°C for 4 hours. After 4 hours the animals were washed three times with M9 buffer with 100ug/mL of ampicillin, 10ug/mL of gentamicin, and 50ug/mL of kanamycin to remove any remaining bacteria. After the last wash, animals were centrifuge to make a pellet that was 80µL in volume. 200µL of an SDS-DTT solution (200mM DTT, 0.25%(w/v) SDS, 20mM HEPES pH 7.3, 3%(w/v) sucrose, pH 8.0) was added to the pellet. The animals were soaked in the SDS-DTT solution for five minutes under gentle agitation to soften the cuticle. After five minutes the animals were washed with egg buffer (25mM HEPES pH 7.3, 118mM NaCl, 48mM KCl, 2mM CaCl_2_, 2mM MgCl_2_) to remove the SDS-DTT solution. 100µL of 15mg/mL pronase E was added to the pellet of soften animals. The solution was pipetted up/down approximately 200 times over 20 minutes at room temperature using a micropipette to digest the cuticle and free the intact cells. Digestion was halted with the addition of 200µL of L-15 insect media + 10% FBS. Cells were pelleted by centrifugation at 9,600 x *g* at 4°C for 5 minutes. The supernatant was removed and the cell pellet was washed again with 1mL of L-15 insect media + 10% FBS for a total of three washed to fully remove the pronase E. The cell suspension was then allowed to rest on ice for 30 minutes to allow the heavier debris to settle on the bottom of the tube. The L-15 insect media containing the suspended cells after resting on ice was filtered through a 40um nylon filter to prevent blockage during cell sorting.

Sorting was performed on a Beckman-Coulter CytoFlex SRT cell sorter with a 100-micron diameter nozzle. The profiles of the GFP expressing cells were compared to cells isolated from wild-type N2 animals to exclude auto-fluorescent cells. Cells were sorted into tubes containing 12.5µL of cell lysis buffer provided by the SMART-seq Ultra Low Input RNA kit (Takara Bio, cat #634888). cDNA was synthesized immediately after cell sorting.

### RNA Sequencing

RNA sequencing libraries were generated using the cDNA synthesized after cell sorting. The Nextra XT DNA Library Preparation kit (Illumina, cat# FC-131-1024) was used to generated libraries for sequencing. Libraries were submitted to the WSU Genomics core for RNA-seq analysis. RNA-seq and related bioinformatic analyses were done following our published protocol (Sellegounder *et al*, 2019).

### RNAi interference

RNAi was conducted using the Ahringer group library and feeding *C. elegans E. coli* strain HT115(DE3) expressing double-stranded RNA (dsRNA) that was homologous to a target gene. (38, 39). Before exposure, all RNAi clone plasmids were isolated, digested with KpnI, and Sanger sequenced using a T7 promoter primer to check for gene specificity. *E. coli* with the appropriate dsRNA vector were grown in LB broth containing ampicillin (100µg/mL) at 37°C for 15∼16 hours and 120µL was plated on modified NGM plates containing 100µg/mL ampicillin and 3mM isopropyl β-D-thiogalactoside (IPTG). The bacteria were allowed to grow for 15∼16 hours at 37°C. Animals were grown on RNAi plates for at least three generations to ensure efficient knockdown of the specific gene. The third generation of animals was synchronized by transferring well-fed second-generation animals to fresh RNAi plates and allowing the animals to lay eggs for 45 minutes at 25°C. After 45 minutes, the adult animals were removed from the plate, and the resulting eggs were allowed to grow at 20°C for 65 hours. *unc-22* RNAi was included as a positive control in all experiments to account for RNAi efficiency.

### Survival assay

Synchronized animals as described above were prepared on RNAi plates and allowed to grow for 65 hours until the animals reached the young adult stage. *Psuedomonas aeruginosa* PA14 bacteria plates were prepared by culturing *P. aeruginosa* PA14 for 15 ∼ 16 hours in Luria broth (LB) at 37°C in a shaking incubator at 200rpm. A 30µL drop of fresh *P. aeruginosa* PA14 culture was placed on the center of a 3.5cm plate of modified NGM media and incubated for 15 ∼16 hours at 37°C. After incubation, the plates were allowed to cool to room temperature and then seeded with synchronized 65-hour old young adult animals. The survival assays were performed at 25°C, and live animals transferred daily to fresh bacterial plates. Animals were scored for survival twice a day and were considered dead when they failed to respond to touch.

### RNA isolation

Gravid adult animals fed specific dsRNA expressing *E. coli* were lysed using a solution of sodium hydroxide and bleach (ration 5:2), washed, and the eggs were synchronized for 22 hours in S-basal liquid media at room temperature. The synchronized L1 larval animals were transferred to modified NGM RNAi plates seeded with specific dsRNA expressing *E. coli* and grown at 20°C for 48 hours until the animals were L4 larva. The 48-hour old animals were then washed with M9 buffer and transferred to modified NGM plates seeded with *E. coli* OP50 and grown for another 24 hours at 20°C to remove any residual dsRNA expressing bacteria. After 24 hours, the animals were collected and washed with M9 buffer and snap frozen in 500µL of Nucleozol (Macherey-Nagel, cat# 740404.200). The total RNA was isolated using the NucleoSpin RNA set for Nucleozol (Macherey-Nagel, cat# 740406.50) following the protocol provided by the manufacture.

### Quantitative real-time PCR (qRT-PCR)

Total RNA was obtained as described above. 500ng of RNA were used to generate cDNA in a 20µL reaction using the New England Biolab’s Lunascript RT supermix kit. The cDNA was diluted prior to qRT-PCR for a final concentration of 5ng/µL. qRT-PCR was conducted by following the provided procotol for New England Biolab’s Luna Universal qPCR Master Mix. 10µL reactions were set up following the manufacturer’s recommendations, and 10ng of cDNA was used per reaction. Relative fold-changes for transcripts were calculated using the comparative *C*_*T*_(2^*-ΔΔCT*^) method and were normalized to pan-actin (*act-1, -3, -4*). Amplification cycle thresholds were determined by the StepOnePlus software. All samples were run in triplicate.

## QUANTIFICATION AND STATISTICAL ANALYSIS

Survival curves were plotted using GraphPad PRISM (version 11) computer software. Survival was considered different from the appropriate control indicated in the main text when *P* <0.05. PRISM uses the product limit or Kaplan-Meier method to calculate survival fractions and the log-rank test, which is equivalent to the Mantel-Haenszel test, to compare survival curves. qRT-PCR results were analyzed using two-sample *t*-tests for independent samples; *P-*values <0.05 are considered significant. All experiments were repeated at least three times, unless otherwise indicated. Statistical details for each figure are listed in its corresponding figure legend.

## Notes

### Competing Interest Statement

The authors have declared no competing interest.

